# Effects of sulfide on the activity of the ammonia-oxidizing archaeon *Nitrosopumilus maritimus* SCM1

**DOI:** 10.64898/2026.05.25.727494

**Authors:** Paula García-Otero, Beate Kraft

## Abstract

Ammonia-oxidizing archaea (AOA) are frequently found in oxygen-depleted marine environments with permanent or temporal presence of sulfide (HS^−^). However, it remains unexplored how sulfide affects the activity of ammonia-oxidizing archaea. We studied the capability of *Nitrosopumilus maritimus* SCM1 to oxidize ammonia when exposed to HS^−^. Ammonia oxidation remained active even after exposure to sulfidic spikes in the lower micromolar range, albeit at reduced rates compared to the absence of HS^−^. However, 90 µM HS^−^ completely inhibited ammonia oxidation. We found no evidence of NO-dismutation under oxygen depletion and presence of HS^−^ (20 µM): the formation of O_2,_ N_2_O and N_2_ did not occur.

All in all, we confirmed ammonia oxidation in *N. maritimus* SCM1 under oxic conditions after sulfide additions, but no evidence of NO-dismutation under sulfidic conditions. Our findings suggest that AOA can recover ammonia-oxidation activity after oxygen re-exposure in regions with periodic sulfide accumulation. However, in permanently sulfidic areas, ammonia oxidation recovery seems unlikely, as NO-dismutation does not appear to be a viable mechanism.

## Introduction

Ammonia-oxidizing archaea (AOA) perform the first step of nitrification, the oxidation of ammonia to nitrite (Schleper & Nicol, 2010; Stahl & De La Torre, 2012). The first step in ammonia-oxidation is catalyzed by the ammonia monooxygenase (AMO) which needs molecular oxygen to function (Dua et al., 1979; Hollocher et al., 1981). Despite their need for oxygen, these ubiquitous microorganisms have been found in several oxygen minimum zones (OMZs) and in anoxic basins, regions with oxygen concentrations below the detection limit (Belmar et al., 2011; Bristow et al., 2016; Cernadas-Martín et al., 2017; Cram et al., 2024; Labrenz et al., 2010; Pitcher et al., 2011; Sollai et al., 2019; Wittenborn et al., 2023). There are various hypotheses that, independently or in combination, could explain their presence in these regions: transient oxygen intrusions delivering oxygen (Buchanan et al., 2024), oxygen originating from the photosynthetic activity of phytoplankton in the second chlorophyll maximum (Garcia-Robledo et al., 2017; Ulloa et al., 2012), or AOA may produce their own oxygen from nitrite through NO-dismutation. This has been shown in isolates (Hernández-Magaña & Kraft, 2024; Kraft et al., 2022) but has yet to be demonstrated in environmental samples. In the proposed pathway nitrite is reduced to nitric oxide by the nitrite reductase (NirK), NO is then dismutated to O_2_ and N_2_O that is further reduced to N_2_ (Kraft et al 2022).

In some of the above-mentioned regions, AOA are also exposed to sulfide (HS^−^): AOA have been frequently found in permanent sulfidic environments such as the Black Sea (Sollai et al., 2019), as well as in systems where HS^−^ accumulates periodically (Berg et al., 2015; Labrenz et al., 2010). Although AOA affiliating with *Nitrosopumilus* were present in sulfidic depths, sometimes at relative high abundances, in all the above-mentioned studies, their abundance generally declined with increasing HS^−^ concentrations. While this trend was also observed in the Black Sea, here AOA still made up more than 10% of the archaeal 16S rRNA reads when HS^−^ exceeded 40 µM. In the Gotland Deep, *Nitrosopumilus* decreased from 20% of relative abundance at oxygen-depleted depths to 5% in the presence of HS^−^ (2 µM) (Berg et al., 2015). A similar decline was reported in the Amberjack Hole (Gulf of Mexico) where *Nitrosopumilus* reached 40% of the total microbial community at oxygen concentrations around 5 µM, and 3% under oxygen-depletion and HS^−^ accumulation (250 µM) (Patin et al., 2021).

*Nitrosopumilus* has also been found in upper and oxic depths of some OMZs that experienced episodes of sulfide plumes with HS^−^ concentrations up to 6 µM (Schunck et al., 2013a) or 30 µM (Ohde & Dadou, 2018). Such sulfidic plumes have been observed in other regions where AOA have been identified (Stewart et al., 2012; Sun & Ward, 2021). However, most of the studies lack simultaneous tracking of HS^−^ and AOA populations. Callbeck et al. (2021) compiled marine areas that experience sulfidic plumes during the austral spring that could potentially expose AOA to HS^−^: the West Indian Shelf (Naqvi et al., 2000), Eastern Tropical South Pacific (Peruvian and Chilean Shelf) (Schunck et al., 2013b; Sommer et al., 2016) and Eastern Tropical South Atlantic (ETSA, Namibian Shelf) (Ohde & Dadou, 2018). *Nitrosopumilus* abundance reached 2% in the Chilean upwelling system, an area that is known for presenting a cryptic sulfur cycle where sulfide and sulfate are rapidly interconverted without net sulfide accumulation (Canfield et al., 2010; Crowe et al., 2018).

The HS^−^ in these OMZs and anoxic basins has two main sources in the water column: it can be released through the activity of sulfate-reducing bacteria (SRB) in sediments (JØrgensen et al., 2010; Scranton et al., 1987) or, in some OMZs, such as the ETSP, it can also be produced by SRB activity in the core of the OMZ (Canfield et al., 2010). In the last scenario, HS^−^ tents not to accumulate since it is consumed by sulfur-oxidizing bacteria (SOB).

Nitrite oxidizers have been shown to be more sensitive to HS^−^ than ammonia oxidizers (Bejarano Ortiz et al., 2013; Bejarano-Ortiz et al., 2015; Bentzen et al., 1995; Erguder et al., 2008; Martienssen et al., 1995). For example the addition of 60 and 100 µM sulfide to sediment slurries reduced or inhibited total nitrification respectively (Joye & Hollibaugh, 1995) and led to nitrite accumulation affecting nitrite oxidizers to a higher degree when compared to AOA and ammonia-oxidizing bacteria (AOB) in estuarine sediments (Caffrey et al., 2007). In sediments, measurable nitrification rates paired with the detection of *amoA genes* at sulfide concentrations between 0.1-0.5mM indicated that AOA could be tolerant to HS^−^ (Caffrey et al., 2007). Similarly, in incubations with water collected from oxygen-depleted depths of the Baltic Sea, sulfidic spikes in the low micromolar range simulating *in situ* conditions, resulted in reduced but persistent nitrification activity as with increasing sulfide concentrations (Berg et al., 2015).

Although field observations suggest that AOA persist in sulfidic environments (Coolen et al., 2007; Sollai et al., 2019, Berg et al., 2015), direct experimental evidence for their physiological response to HS^−^ remains scarce.

Additionally, archaeal *amoA* have been retrieved from sludge reactors after sulfide additions at mM concentrations suggesting a potential tolerance to HS^−^ (Erguder et al., 2008). However, to our knowledge, no study has directly explored the effect of HS^−^ on the physiology of AOA in controlled experiments. For AOB, Sears et al. (2004) showed that nitrification rates recovered after HS^−^ exposure under aerated conditions. In that study, ammonia oxidation ceased while sulfide and oxygen were present, and resumed after all the sulfide was oxidized. Whether AOA respond similarly remains unknown.

Since NO-dismutation was shown to occur in *N. maritimus* SCM1 (Kraft et al., 2022), one possible explanation for their survival in sulfidic environments is that ammonia-oxidation is sustained through internal oxygen production. However, this would require tight coupling of oxygen production and consumption.

In this study, we selected the most studied marine AOA, *Nitrosopumilus maritimus* SCM1, to assess its ability to sustain ammonia oxidation at low oxygen and sulfidic concentrations, and if *N. maritimus* SCM1 can perform NO-dismutation under oxygen-depleted and sulfidic conditions.

## Materials and methods

*N. maritimus* SCM1 was kindly provided by Martin Könneke. It was grown under oxic conditions in HEPES-buffered synthetic Crenarchaeota medium (pH 7.5) (Martens-Habbena & Stahl, 2011) with 6 mM of sodium bicarbonate (Kraft et al., 2022) and 1 mM of ammonium. Growth was monitored by ammonia consumption (Bower & Holm-Hansen, 1980). Purity of cultures was assessed prior to the start of the experiment via DAPI staining and microscopy.

### Ammonia oxidation under reduced oxygen concentrations

The purpose of this experiment was to examine if *N. maritimus* SCM1 can perform ammonia oxidation when exposed to sulfide and when oxygen is present at low micromolar concentrations. Two sets of incubations were performed in triplicates: In the first set (experiment 1A) sulfide was added at concentrations of 0, 10 and 25 µM and killed controls were performed in parallel. In the second set (experiment 1B), sulfide was added at concentrations of 0, 8, 3, 86 µM, however without killed controls.

The experiment was started when the preculture had consumed the ammonium down to a concentration of 50 µM. Prior to the experiment, the culture was flushed with argon (99.99%) for 45min and 170 mL were immediately transferred to serum bottles of 250mL, leaving therefore an initial headspace of 80 mL (Figure 1). The bottles were closed with blue butyl rubber stoppers (Chemglass) and metallic crimps and flushed for 15 more min with Argon to completely deplete the oxygen. The experiment was conducted at constant room temperature (22 °C) in the dark. The bottles were mixed by constant stirring at 250rpm. Oxygen was monitored every 20s with trace luminescence oxygen sensors (hereafter, optodes), glued in the lower part of the bottle (Lehner 2015). Samples were collected at 0, 4, 6, 20, 27, 45, 51, 69, 75, and 92 h. Sulfide was added 4 h after the beginning of the experiment, followed by additions of oxygen to ensure oxygen concentrations stayed above 1 µM. Spikes of oxygen were added during the experiment when oxygen concentrations decreased (see Table S1). All substrate additions and sampling were performed with gastight glass syringes (Hamilton, Switzerland) to avoid oxygen contamination. For each sample the same volume of sterile air or argon was introduced to ensure the same pressure conditions along the incubation. Samples for ammonium and sulfide measurements were collected in Eppendorf tubes at all sampling points. Sulfide samples were preserved with 50 µL of Zn acetate (5%). For the killed-controls autoclaved *N. maritimus* culture was used and sulfide concentrations were adjusted to 10 and 25 µM.

**Figure 1.**
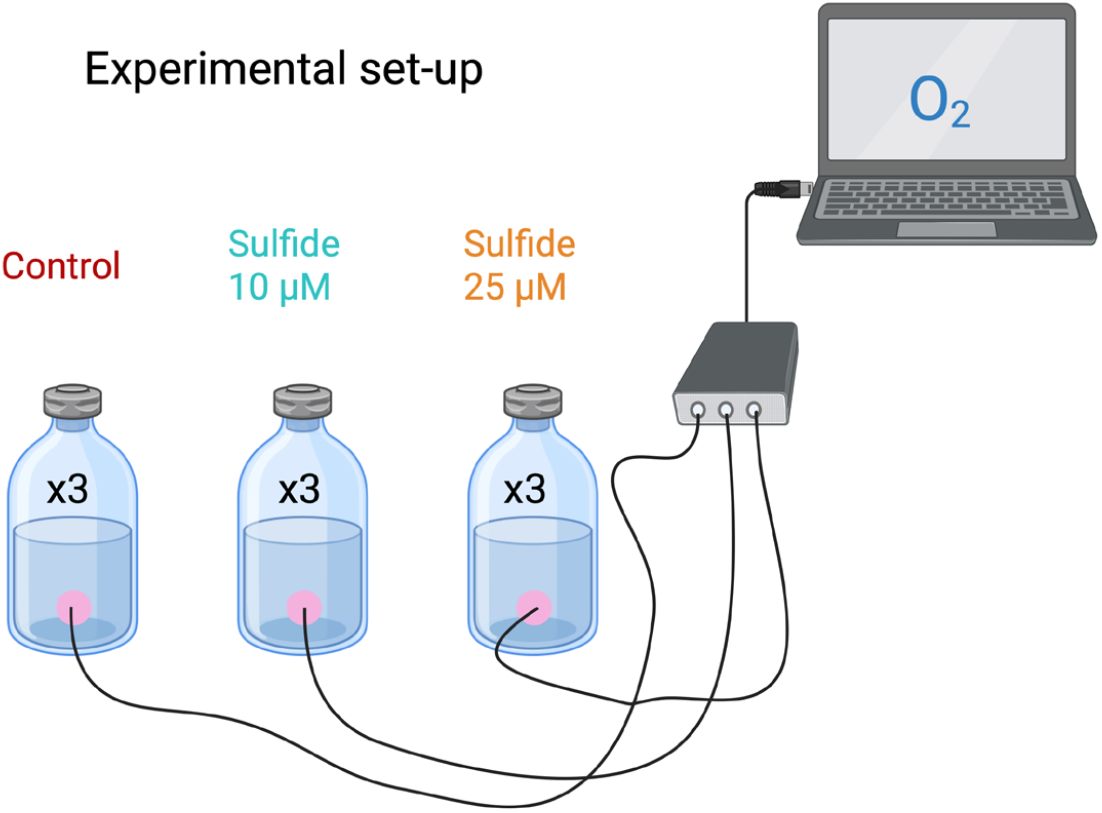
Experimental setup of the incubations. Serum bottles with optodes (pink) with different sulfide concentrations with continuous oxygen monitoring.

### NO-dismutation under oxygen depletion and sulfide exposure

An aerobic culture of *N. maritimus* was grown with 1mM of ^15^NH_4_^+^ until it was completely depleted and therefore converted to ^15^NO_2_^−^. The culture was purged with argon (99.99%) for 45 min and immediately transferred to custom-made capillary glass bottles of 300 mL through a custom-made set-up (Tiano et al 2014, Kraft et al 2022). The following incubations were performed in triplicates respectively: oxygen-depleted incubations, oxygen-depleted incubations with 20 µM HS^−^ and oxygen-depleted killed and abiotic controls. For the killed control 3 ml of Argon-flushed, saturated HgCl_2_ solution was added through the capillary. Oxygen and NO were monitored with optodes (glued to the inside of the bottle) (Lehner et al., 2015) and NO-microsensors (Unisense, Denmark) inserted through sensor ports (Tiano et al., 2014) respectively. The bottles were incubated in a water bath at 28 °C, in the dark, and with constant stirring with glass coated stirring bars (VWR, UK) at 250 rpm. Every time a sample was collected, the same volume was replaced with anoxic and sterile medium. Samples were collected at 20, 25, 30, 35, 42, 51, 67, 75, and 93 h with gastight Hamilton syringes connected to stainless steel needles (Ochs, Germany). Samples were transferred to 3 mL Exetainers (Labco, UK) free of headspace prefilled with 50 µL of saturated HgCl_2_ and analyzed by coupled gas chromatography-isotope ratio mass spectrometry (GC-IRMS) (Delta V Plus isotope ratio mass spectrometer, Thermo) (Dalsgaard et al., 2013) to evaluate ^30^N_2_ and ^46^N_2_O production. O_2_ concentrations were corrected for the interference of NO according to (Kraft et al., 2022). Oxygen optodes and NO sensors were calibrated before the incubations. Despite minimizing oxygen contamination as much as possible during sample collection, a small intrusion of oxygen always occurred. As a result, abrupt peaks in the oxygen plots mark the different sampling points.

### Chemical analyses

Ammonium (Bower & Holm-Hansen, 1980) and sulfide (Cline, 1969) were analyzed spectrophotometrically. Laboratory tests of Crenarchaeota media showed that high nitrite concentrations accumulating in the media during growth of *N. maritimus* interfered with sulfide measurements. Therefore, sulfide samples (4 mL, fixed with Zinc acetate) were filtered through Whatman™ Grade GF glass microfiber filters (1.6 µm pore size). Thereafter, the filters were submerged in in 2-3 mL of MQ water, followed by mixing and addition of cline reagent. 1ml of was transferred to an Eppendorf tube and centrifugated at 15.000 g to make sure that any residual parts of the filter settled to the bottom of the tube and measured.

Ammonia oxidation rates were calculated and compared between the different sulfide treatments. We applied a linear regression model to the ammonia consumption rates once ammonia started to be consumed and did an ANOVA to study if there was a statistically significant difference between the slopes of the different sulfide incubations. Afterwards, we performed a pairwise t-test with Bonferroni correction to assess if the slopes of the ammonia consumption were statistically different from each other.

## Results

### Effect of sulfide on ammonia oxidation

To investigate the ammonia oxidation under the presence of oxygen and sulfide, *N. maritimus* was exposed to different HS^−^ concentrations (0, 10, and 25 µM). HS^−^ was added 4h after the start of the experiment. As oxygen was abiotically reduced by the HS^−^ it was added back to the headspace, ensuring that oxygen concentrations always stayed above 1 µM (Figure S1). Oxygen was just below 1 µM for few hours after sulfide was completely oxidized.

Ammonia oxidation occurred in all the incubation in the first 4 h of the experiment prior to the sulfidic spike. During the presence of sulfide, no ammonia oxidation was detected. After HS^−^ was oxidized, ammonia oxidation started again. Ammonia consumption was statistically different between the treatment without sulfide and 10 and 25 µM (p-value<0.05). However, the ammonia consumption rates showed no statistically significant difference (p-value>0.05) across the different sulfide treatments (Table S2). Ammonia oxidation in the non-sulfidic control was 1.36±0.04 µM/h. This rate was reduced by 69±1% and 72±5% with 10 and 25 µM of sulfide respectively. In the incubation with 25 µM HS^−^, the culture needed 17 h after HS^−^ oxidation to restart ammonia oxidation (Figure 2A). No ammonia was consumed in any of the abiotic controls, therefore confirming biotic ammonia oxidation by *N. maritimus* SCM1.

**Figure 2.**
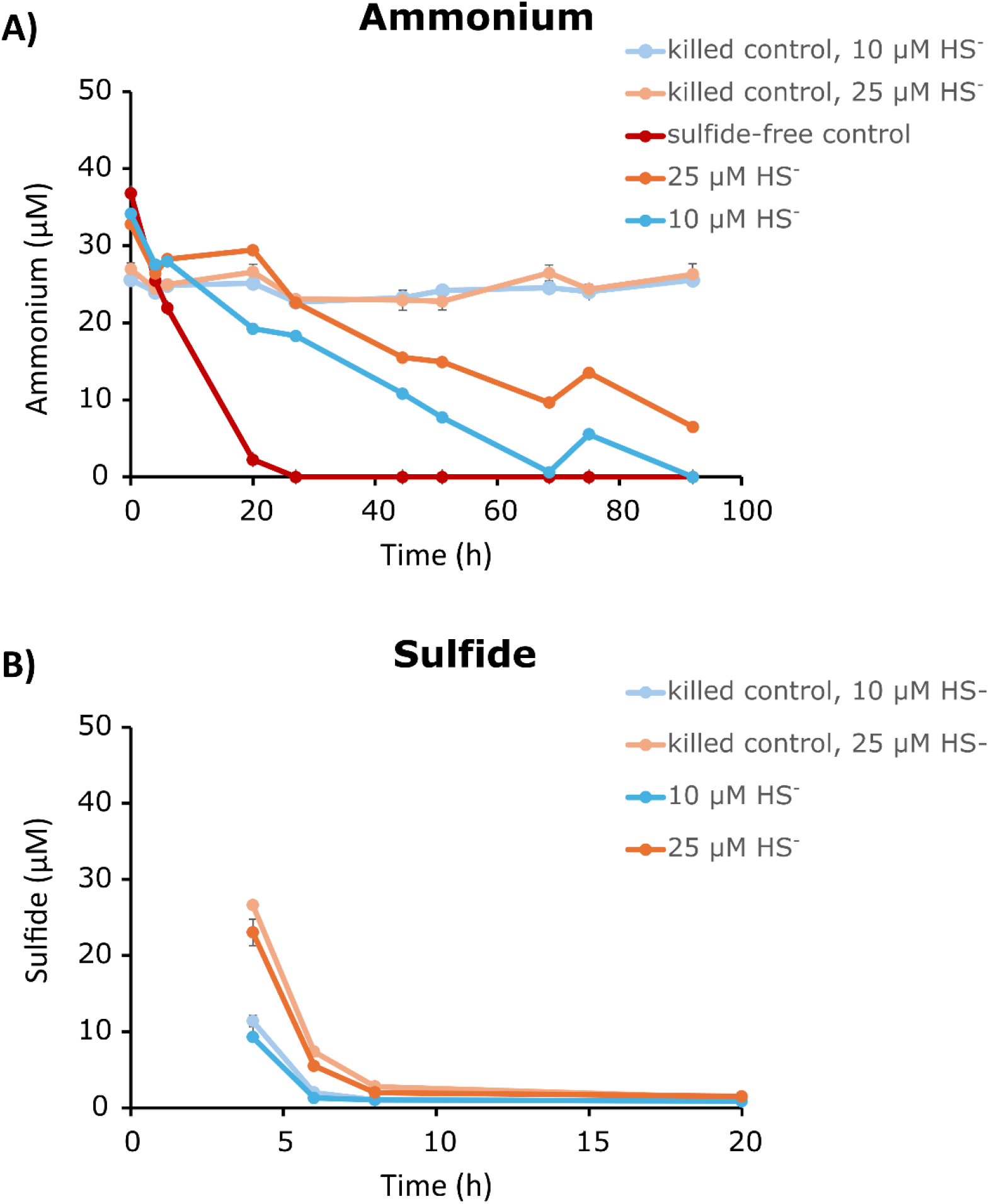
Ammonium (A) and sulfide (B) concentrations in oxic incubations of *Nitrosopumilus maritimus* SCM1 exposed to sulfide. Datapoints show an average of the triplicates with error bars indicating the standard deviation. Incubations represented as following: control without sulfide (red), 10 µM HS^−^ (dark blue), 25 µM HS^−^ (orange), abiotic control with 10 µM HS^−^ (light blue), abiotic control with 25 µM HS^−^ (light orange). Some standard deviations are smaller than the symbols and not visible.

After the sulfidic spike at 4h, the initial HS^−^ concentrations were 9 ±3 and 23 ± 2 µM. All the sulfide in abiotic and biotic controls was consumed in 2 and 4 h after the sulfide spike in both 10 µM and 25 µM HS^−^ incubation bottles respectively. No statistically significant differences were found between the killed controls and abiotic controls (p-value>0.05) when comparing 10, and 25 µM incubations. The similarity of the HS^−^ oxidation rates between killed controls and biotic controls indicates abiotic sulfide oxidation (Figure 2B).

All incubations were oxic. However, upon complete oxidation of HS^−^, oxygen increased sharply and earlier in the bottles with 10 µM of H_2_S compared to those with 25 µM (Figure S1). The same pattern was also observed in the killed controls (Figure S1).

In a second experiment, the effect of broader range of sulfide concentrations was tested. HS^−^ (8.3 ± 0.8, 33.5 ± 3.1 and 85.8 ± 4.2 µM) was added at the beginning of the experiment, reducing the oxygen concentrations to 0.5-1µM in the first 30 min. Here, similar trends were observed as in the first experiment: Sulfide additions caused ammonia oxidation to completely cease for at least 28h.

While in the incubation with initial 34 µM HS^−^ ammonia oxidation was still observed albeit at a very slowed down rate (inhibition of 67 ± 5%), ammonia oxidation was completely inhibited after exposure to 86 µM of sulfide (ammonia oxidation inhibition of 97% ± 2%). Ammonia consumption rates (Figure S2) were statistically different between all the different sulfidic incubations (p-value <0.05) and compared to the control except for 34 µM and 86 µM that presented no statistical difference (p-value >0.05) (Table S2). The ammonia consumption rate in the non-sulfidic control was 1.44 ± 0.06 µM/h, comparable to the first experiment. After all sulfide was oxidized dissolved oxygen concentrations increased again in a similar way as was observed in the first experiment.

### Activity of *N. maritimus* exposed to sulfide under oxygen-depleted conditions

Oxygen-depleted incubations with *N. maritimus* SCM1 were amended with 20 µM of sulfide to test the effect of HS^−^ on NO-dismutation. These incubations were compared to incubations without sulfide and killed controls.

Upon oxygen consumption in the first two hours, oxygen production started reaching up to 200 nM in all incubations (Figure 3). After 20 h of oxygen production, 20 µM of HS^−^ was added to the sulfidic incubations. In the sulfide-free incubations, oxygen production and consumption were observed displaying patterns similar to those reported in previous experiments (Kraft et al., 2022). During sampling, small oxygen intrusions were introduced indicated with black arrows in Figure 3. After the addition of sulfide, oxygen production ceased. Oxygen introduced during sampling was consumed.

**Figure 3.**
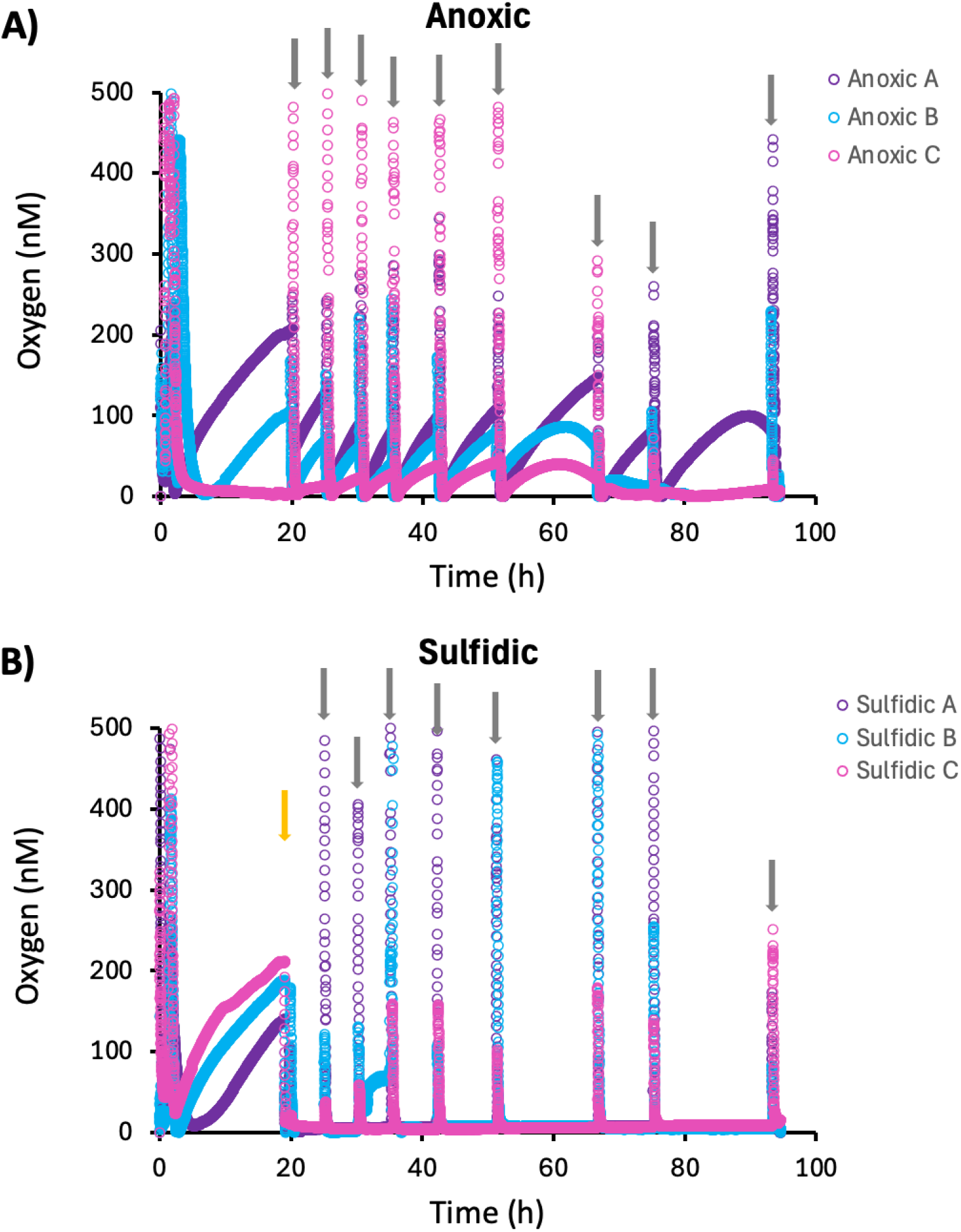
Oxygen concentrations in the oxygen-depleted incubations of *Nitrosopumilus maritimus* SCM1 in the non-sulfidic control (A) and sulfidic incubation (B). A, B and C represent parallel replicates of the incubations. Black arrows indicate the sampling points where small oxygen intrusions occurred, and the orange arrow indicates the sulfide spike.

In the sulfide-free control, nitric oxide accumulation was coupled to oxygen production as reported previously (Kraft et al 2022) and NO concentrations reached a maximum of 1-4 µM NO along in the first 20 h of the incubations (Figure S3). All sulfidic replicates showed NO accumulation prior to sulfide additions. After the sulfidic spike, NO accumulation stopped.

The initial measured sulfide concentration was 14±1.5 µM and decreased to 5 ±1 µM by 75 h (Figure S4).

The sulfide present throughout the incubation would prevent oxygen from accumulating even if it was produced via NO-dismutation. In order to track any potential NO-dismutation activity incubations contained ^15^NO_2_^−^. If NO-dismutation occurred, the ^15^NO_2_^−^ present in the culture, would be converted to ^46^N_2_O and ^30^N_2_. While the oxygen-depleted bottles showed ^46^N_2_O and ^30^N_2_ accumulation up to 282 ± 296 nM and 3903 ± 2717 nM respectively after 93 h, the sulfidic bottles did not show accumulation (Figure 4). Just in one replicate of the sulfidic incubations, N_2_O production was detected in the last time point at 93 h.

**Figure 4.**
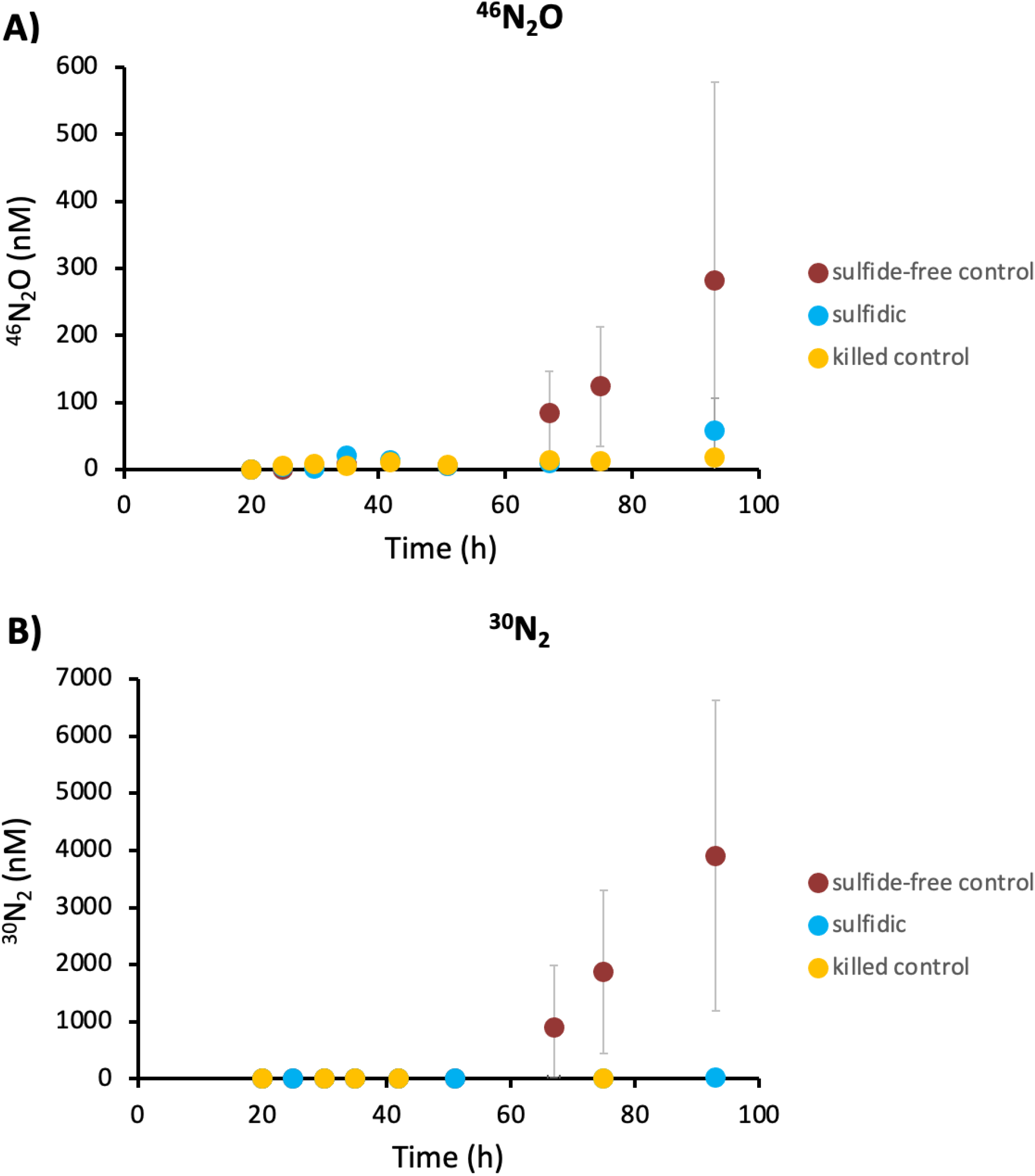
Nitrous oxide (A) and nitrogen (B) accumulation from incubations of *N. maritimus* SCM1 with ^15^N-nitrite under oxygen depletion. Datapoints show an average of triplicates, error bars represent the standard deviation: oxygen depleted incubations (red), sulfidic (blue) and abiotic control (orange). Some standard deviations are smaller than the symbols and not visible.

## Discussion

### Ammonia oxidation follows abiotic sulfide oxidation

Recovery of ammonia oxidation activity was immediate after abiotic HS^−^ depletion. Ammonia oxidation was delayed proportionally to the initial HS^−^ concentrations the AOA were exposed to, ammonia oxidation recovered after exposure of up to 34 µM, and 86 µM HS^−^ completely inhibited it. This observed tolerance is higher than reported inhibitory concentrations for AOA communities from the Gotland Deep (4.1–16.6 µM; Berg et al., 2015) and AOB enrichments (7 µM; Sears et al., 2004), but comparable to inhibition observed in estuarine sediments (60–100 µM; Joye & Hollibaugh, 1995) and aerated sludge (~90 µM; Bejarano Ortiz et al., 2013), though the latter studies did not distinguish between AOA and AOB or consistently monitored oxygen.

In our experiments ammonia oxidation recovered after sulfide was abiotically oxidized. Sears et al. (2004) showed similar trends in enrichments with AOB related to *Nitrosomonas:* aerated conditions after a continuous pumping of 60mM of HS^−^ showed a recovery of ammonia oxidation activity after HS^−^ was fully oxidized, but at slower rates when compared to pre-sulfidic exposure. Nevertheless, this observation in comparison with this study suggest that certain AOB can tolerate sulfide at concentrations that are several magnitudes higher than AOA.

Oxygen intrusions have been observed in sulfidic areas of the Baltic Sea (Beier et al., 2019) that could be compatible with the nature of our experiment, where ammonia oxidation just occurs after sulfide depletion caused by oxygen. This could imply that ammonia oxidation by AOA could be sustained by small periodic oxygen intrusions in sulfidic basins. The same authors showed that *amoA/B/C* and *nirK* genes were increasingly overexpressed in time-series incubations under different conditions: oxic and mixed oxic-sulfidic (10 to 0.2 µM HS^−^ from the start to end of the experiment). However, the expression remained constant but high in the sulfidic incubations (25 to 7 µM HS^−^ from initial to end of the incubation). No oxygen was monitored during these incubations.

Other sulfur-containing molecules have been shown to inhibit *amoA* expression in environmental samples. Dimethyl sulfide (DMS) inhibited ammonia oxidation and *amoA* expression in activated sludge with AOB related to *Nitrosomonas europaea*, but the ammonia oxidation inhibition differed among populations (Fukushima et al., 2014). Carbon disulfide (CS_2_) reduced ammonia oxidation in *Nitrosomonas europaea*, but it did not completely inhibit it. It was proposed that CS_2_ might react with the amino acids close to the copper active site (Hyman et al., 1990).

Inhibition of ammonia oxidation by HS^−^ using sludge (without making a distinction between AOA and AOB) showed that ammonia oxidation was non-competitive (Bejarano-Ortiz et al., 2015). Our experiments did not show ammonia oxidation until HS^−^ was completely oxidized. When oxygen was present, HS^−^ seemed to inhibit ammonia oxidation in a reversible way up to a concentration of 86 µM that produced an irreversible inhibition. We cannot distinguish if the inhibition was competitive, non-competitive or mixed, since ammonia-oxidation did not occur in presence of sulfide. To discard a competitive inhibition, the experiment would need to be performed with higher ammonia concentrations compared to sulfide. Independently of the mechanism of ammonia inhibition, it was shown that HS^−^ affects ammonia oxidation in *N. maritimus*.

Abiotic HS^−^ consumption in the killed controls indicate that most of the HS^−^ was oxidized by the oxygen present in the incubations, but some soluble cations of the media components could also have contributed to the precipitation of HS^−^ (Sears et al., 2004). They hypothesized S^2-^ could have reacted with copper and therefore scavenged it from the AMO enzyme, reducing therefore their activity. Both AMO and NirK have copper in their active center.

Oxygen accumulated in all incubations with oxygen additions following complete HS^−^ oxidation, with higher HS^−^ concentrations requiring longer to reach this point. We attribute this to equilibration between the liquid phase and headspace: once HS^−^ was fully oxidized, the culture no longer acted as an oxygen sink, and dissolved oxygen equilibrated with the excess oxygen in the headspace (Table S1).

### Sulfide affects ammonia oxidation rates and NO-dismutation

HS^−^ (20 µM) completely inhibited NO-dismutation in *N. maritimus* SCM1. The fact that O_2_ did not accumulate could be explained by abiotic reaction of oxygen with HS^−^. If NO-dismutation was occurring, even though O_2_ would not accumulate, N_2_O (and potentially N_2_) would be produced according to the proposed pathway (Kraft et al., 2022). Stoichiometrically, we would expect twice as many molecules of N_2_O as of O_2_ and we did not observe any accumulation.

NO accumulation ceased after HS^−^ additions, suggesting that *NirK* could be affected by HS^−^. In sulfidic waters of the ETSP, *NirK* (involved in the reduction of nitrite to NO) and nitrous oxide reductase (*nosZ)*, both with copper active sites (Godden et al., 1991; Nojiri et al., 2007), have a lower expression when compared with the homologous Fe-nitrite reductase NirS (Schunck et al., 2013b, Léniz et al., 2017). On the other hand, NO may have been scavenged by sulfide.

It is very well possible that NO-dismutation is not a feasible metabolic pathway under sulfidic conditions in the environment. This could be the case in sulfidic depths of many permanent anoxic basins such as the Black Sea where the water column is permanently euxinic and AOA abundance significantly dropped with increasing sulfide concentrations (Sollai et al., 2019). However, it remains unknown if AOA could switch between NO-dismutation under oxygen depletion and absence of sulfide and ammonia oxidation in the presence of low concentrations of oxygen and sulfide. This could be the case in some OMZ exposed to sulfide plumes or anoxic basins with periodic sulfidic intrusions (Beier et al., 2019; Berg et al., 2015). On the other hand, ammonia oxidation could be sustained with small oxygen intrusions in the euxinic areas of the Baltic Sea.

NO-dismutation (Hernández-Magaña et al., 2023; Kraft et al., 2022) serves as a possible explanation to explain the survival of AOA in oxygen-depleted waters. However, this study suggests that this mechanism cannot be sustained in sulfidic environments. We cannot completely rule out the possibility of AOA performing NO-dismutation in sulfidic environments, since SOB could be utilizing the HS^−^ near the AOA cells and therefore, reducing its exposure to HS- (Erguder et al., 2008; Park et al., 2014; Sollai et al., 2019). For example, in sulfide-rich mangrove swamps, AOA and SOB have been shown to be physically associated (Muller et al., 2010).

## Conclusion

This study provides direct experimental evidence of how sulfide affects AOA physiology. *N. maritimus* SCM1 oxidized ammonia at low micromolar oxygen concentrations after exposure to sulfide but the degree of inhibition of ammonia oxidation rates increased with increasing HS^−^ concentrations. Additionally, NO-dismutation was not observed under sulfidic and anoxic conditions. Together these two experiments provide new insights into the physiology and ecology of environmental *Nitrosopumilus* exposed to sulfidic conditions.

These results suggest that AOA can resume ammonia oxidation following transient sulfidic events where oxygen drives sulfide oxidation, as observed in the Baltic Sea or OMZs experiencing episodic sulfide plumes. However, NO-dismutation appears an unlikely survival mechanism in permanently euxinic environments, leaving the persistence of AOA at sulfidic depths an open question.

## Acknowledgements

We thank Sophie Lorenz for help in optimizing HS^−^ measurements. This work was supported by the Villum Foundation, Denmark (Villum Young Investigator Grant 00025491 and 00060799 to B.K).

## Author contribution

PGO and BK contributed to the experimental design. PGO performed the experiments and the data analysis with input from BK. PGO and BK wrote the manuscript.

## Notes

### Competing Interest Statement

The authors have declared no competing interest.

