## Supplementary information for "Effects of sulfide on the activity of the ammonia-oxidizing archaeon *Nitrosopumilus maritimus* SCM1"

### Supplementary Material

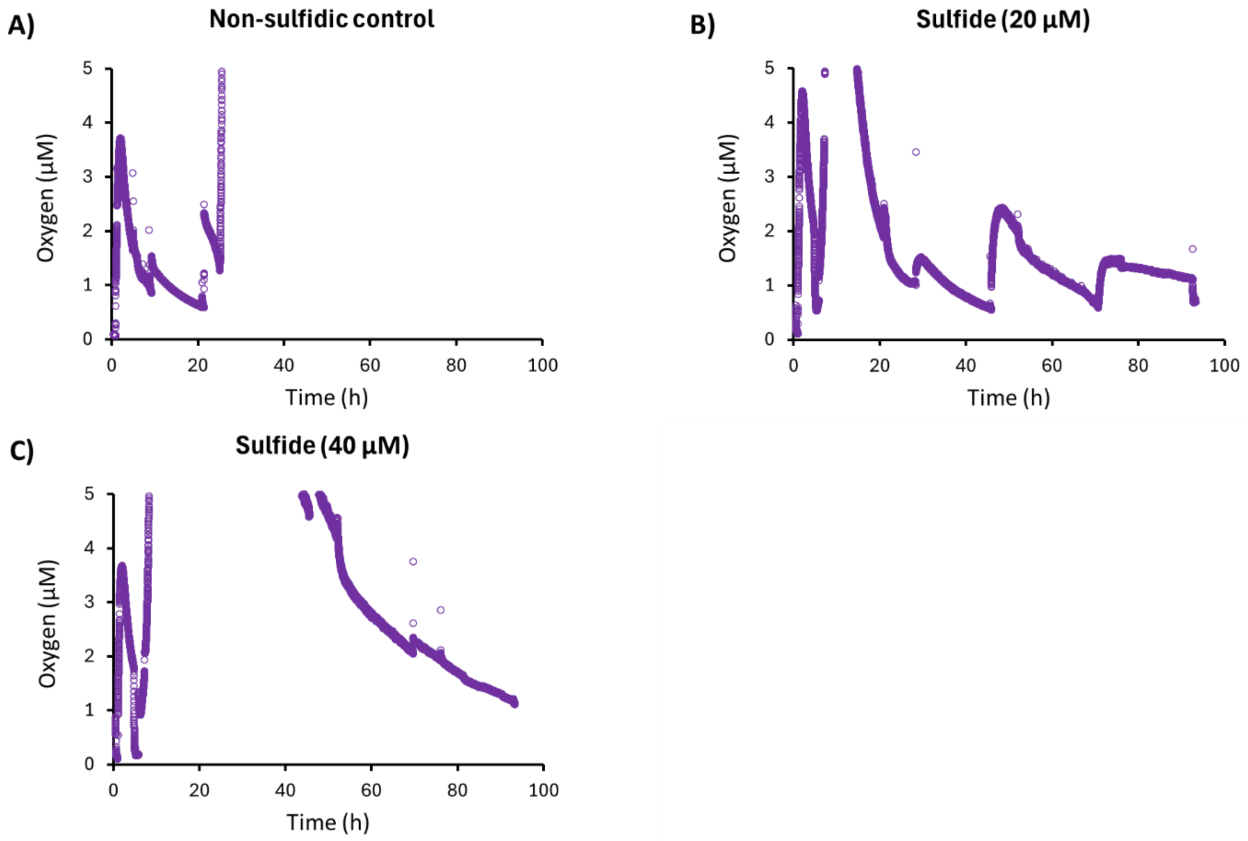


**Figure S1**. Examples of the oxygen concentrations in experiment 1A in one of the replicates. The killed controls are not shown since they showed oxygen concentrations above 5µM along the incubations, therefore out of the measurement range.

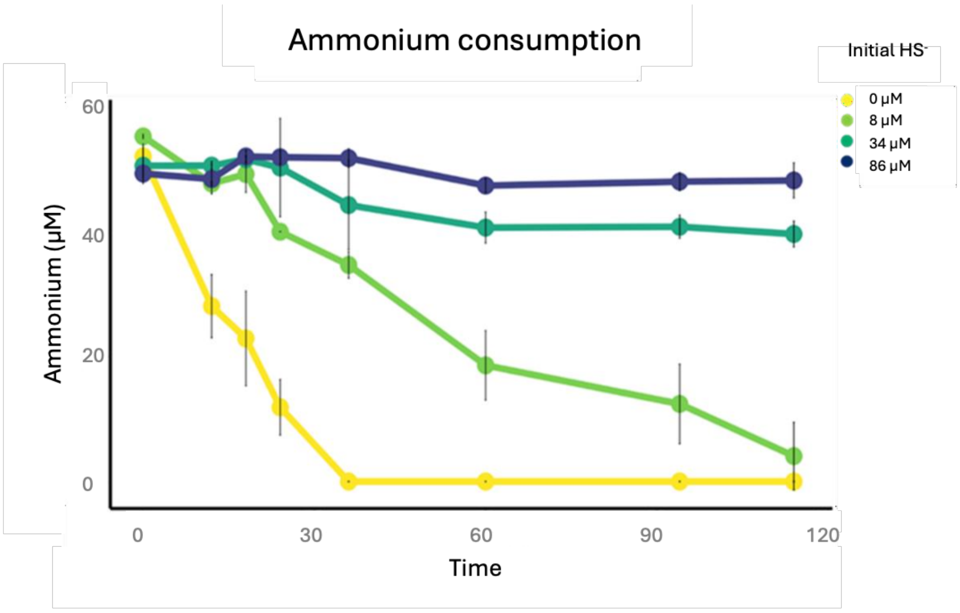


**Figure S2.** Ammonium concentration in oxic incubations of *Nitrosopumilus maritimus* SCM1 exposed to different initial sulfide concentrations: 0 µM (yellow), 10 µM (light green), 35 µM (dark green), 85 µM (blue). The error bars represent the standard deviation of the triplicates.

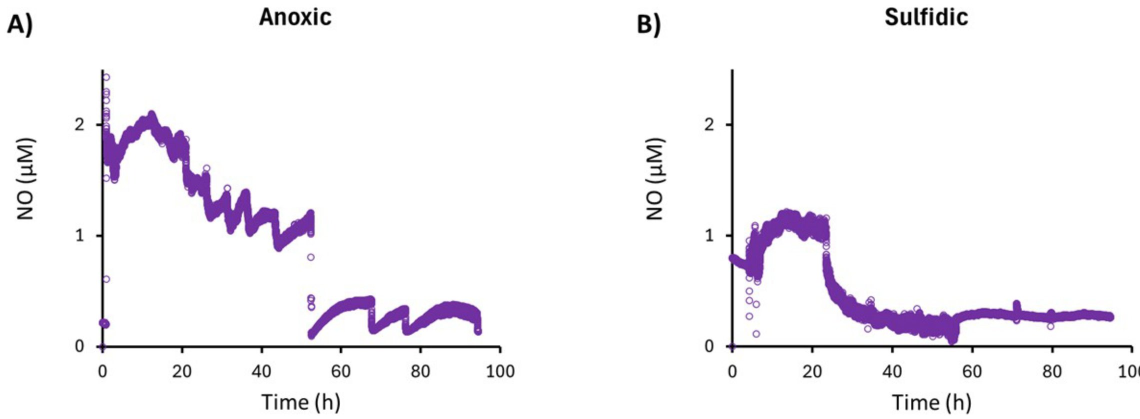


**Figure S3.** Examples of nitric oxide (NO) concentrations (in µM) along the anoxic incubations under anoxic non-sulfidic conditions (A) and sulfidic conditions (B).

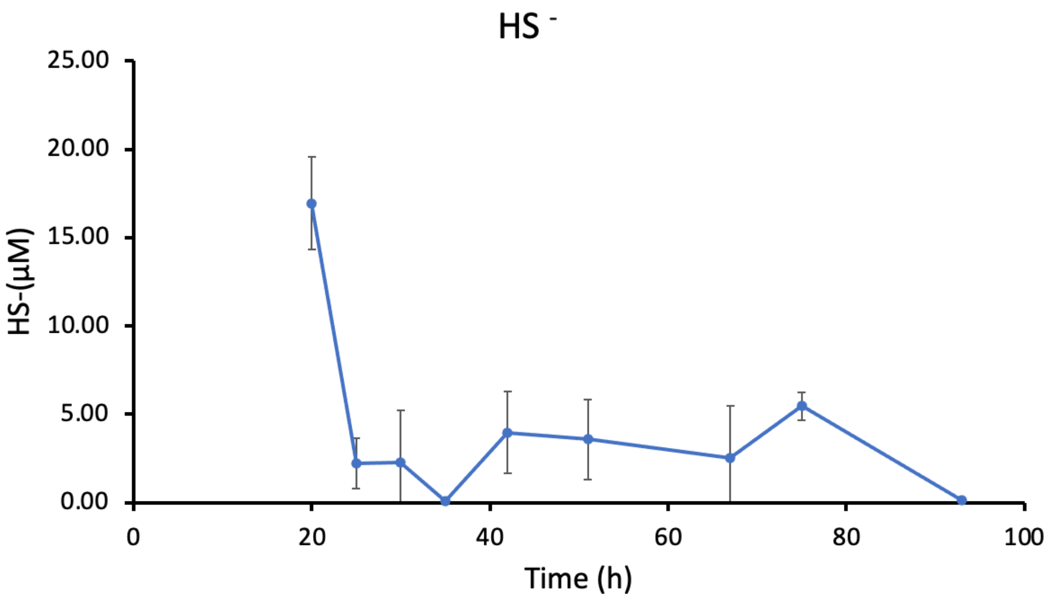


**Figure S4.** Sulfide concentration (µM) along the anoxic incubation. The error bars represent the standard deviation of the triplicates.

**Table S1**. Theoretically expected maximal abiotic sulfide oxidation capacity for the total amount of oxygen that was added during the experiment. In the table, the air (in mL) injected between the t0-t2 in the serum bottles is indicated and the corresponding calculated $O_2$ concentrations. Ideal gases were assumed for the calculations.

| **Incubation (µM of $HS^-$)** | **Air (mL) until t2** | **$O_2$ (ml)** | **$O_2$ (µM)** | **Theoretical $HS^-$ abiotic Oxidation** |
|---|---|---|---|---|
| 10 µM | 3 | 0.63 | 103.8 | 51.9 |
| 25 µM | 3.5 | 0.74 | 121.1 | 60.6 |
| Killed control, 10 µM | 3 | 0.63 | 103.8 | 51.9 |
| Killed control, 25 µM | 2.5 | 0.53 | 86.5 | 46.3 |

**Table S2.** P-values corresponding to the Bonferroni pairwise t-test results corresponding to 1) ammonia oxidation in experiment 1A (1a), experiment 1B (1b), and 2) sulfide oxidation. Bold letters indicate the initial sulfide concentration in the different incubations. K represents the killed controls. P-values <0.05 indicate a difference between the different incubations.

**1) Ammonia oxidation**

| **1a) Experiment 1A** | **0** | **10** |
|---|---|---|
| **10** | 3.70E-06 | - |
| **25** | 2.90E-06 | 1 |

| **1b) Experiment 1B** | **0** | **86** | **8** |
|---|---|---|---|
| **86** | 2.10E-08 | - | - |
| **8** | 3.80E-07 | 2.00E-04 | - |
| **34** | 3.30E-08 | 1 | 7.40E-04 |

| **2) $HS^-$ oxidation experiment 1A** | **10** | **10k** | **25** |
|---|---|---|---|
| **10k** | 1 | - | - |
| **25** | 4.60E-02 | 1 | - |
| **25k** | 6.80E-02 | 4.12E-01 | 1 |
